# Uni-Fold MuSSe: De Novo Protein Complex Prediction with Protein Language Models

**DOI:** 10.1101/2023.02.14.528571

**Authors:** Jinhua Zhu, Zhenyu He, Ziyao Li, Guolin Ke, Linfeng Zhang

## Abstract

Accurately solving the structures of protein complexes is crucial for understanding and further modifying biological activities. Recent success of AlphaFold and its variants shows that deep learning models are capable of accurately predicting protein complex structures, yet with the painstaking effort of homology search and pairing. To bypass this need, we present Uni-Fold MuSSe (Multimer with Single Sequence inputs), which predicts protein complex structures from their primary sequences with the aid of pre-trained protein language models. Specifically, we built protein complex prediction models based on the protein sequence representations of ESM-2, a large protein language model with 3 billion parameters. In order to adapt the language model to inter-protein evolutionary patterns, we slightly modified and further pre-trained the language model on groups of protein sequences with known interactions. Our results highlight the potential of protein language models for complex prediction and suggest room for improvements.

## 1 Introduction

Proteins as well as their interactions substantiate a majority of biological activities. Therefore, accurately solving the structures of protein complexes provides key insights in understanding and further modifying these activities. A preliminary to protein complex prediction is to solve the structures of monomeric proteins. Thanks to the success of AlphaFold (Jumper et al., 2021) as well as its subsequent works (Mirdita et al., 2022; Li et al., 2022), *de novo* prediction of monomeric protein structures with atomic-level accuracy is now a regular tool in protein analysis. Based on AlphaFold, AlphaFold-Multimer (Evans et al., 2022) further showed that with inter-chain evolutionary information, retrained AlphaFold models are capable of accurately predicting protein complex structures.

Overall, these AlphaFold-based models rely on the evolutionary patterns embodied by the multiple sequence alignments (MSAs) to make accurate predictions. Specifically in AlphaFold-Multimer, a heuristic pairing strategy is introduced, which greedily matches the homologous sequences of target protein monomers by species similarity. However, obtaining these MSAs requires painstaking work of sequence searching in very large databases, which becomes the temporal bottleneck of the prediction pipeline. Also, the models cannot be well generalized to targets with few homologous sequences. To address these issues, an alternative approach was proposed, which replaced the MSAs with sequence representations from protein language models (PLM) (Rao et al., 2019; Elnaggar et al., 2020; Rives et al., 2021), in the hope of learning the evolutionary patterns directly from primary sequences (Lin et al., 2022; Wu et al., 2022). These PLMs were pre-trained to predict randomly masked tokens similarly to natural language models (Devlin et al., 2018), but on extensive protein sequence corpora with the vocabulary of amino acid types. In general, these models obtain approaching or equivalent accuracies on monomer prediction tasks compared with AlphaFold, while bypassing the effort of homology searching.

Nevertheless, the use of PLMs in protein folding is so far limited to monomers. Although some tricks were proposed to predict complex structures with ESMFold (Lin et al., 2022), such as concatenating protein chains with gaps or soft linkers, the accuracies were much limited. A specific challenge of using PLMs for protein complex prediction is that all current PLMs were trained with individual protein chains, leaving it in doubt whether the sequence representations can capture inter-chain evolutionary patterns and further guide the complex prediction task. To answer this question, we i) trained a protein complex prediction model using the representations of input primary sequences from an existing PLM, ESM-2 (Rives et al., 2021) with 3 billion parameters; ii) enhanced the PLM with chain relative positional encoding, and further pre-trained it with the corpora of protein interactions; and iii) check if the further pre-training improves the folding accuracy.

To this end, we present Uni-Fold MuSSe (<u>Mu</u>ltimer with <u>S</u>ingle <u>Se</u>quence inputs), an end-to-end model which *de novo* predicts protein complex structures from primary sequences only. We found that by retraining a multimeric folding model, the representations from ESM-2 were applicable to complex prediction with satisfying accuracy, largely outperforming existing PLM-based baselines, despite a gap to MSA-based models represented by AlphaFold-Multimer. By further pre-training the PLMs on protein interaction corpora, the retrained complex prediction model, Uni-Fold MuSSe+, displayed a better performance. Also, on several test cases that AlphaFold-Multimer failed to work, Uni-Fold MuSSe+ obtained correct predictions.

## 2 Related Work

### Protein Structure Prediction

*In silico* protein structure prediction aims to predict protein structures from the amino acid sequences purely with computational methods. AlphaFold (Jumper et al., 2021) is the first deep learning model that claimed to achieve “atomic-level” accuracy, based on which AlphaFold-Multimer (Evans et al., 2022) achieved high accuracy on protein complex prediction. To facilitate the open-source community of deep protein folding models, Uni-Fold (Li et al., 2022) re-implemented both AlphaFold and AlphaFold-Multimer in PyTorch with equivalent accuracy, and released the training code in the repository. In this paper, we built the multimeric folding model following the implementation of Uni-Fold Multimer.

### Protein Language Models

Inspired by the success of pre-trained language models (Devlin et al., 2018; Radford et al., 2018), encoding the amino sequences of proteins with large language models has been widely explored. Generally, protein language models (PLMs) are trained on large protein sequence corpora to predict randomly masked amino acids in the sequences, whose representations are leveraged in downstream tasks. Early works (Rives et al., 2021; Elnaggar et al., 2020) showed the effectiveness of using PLMs in secondary structure prediction and contact prediction. Later, these PLMs were improved with more curated architectures and larger parameters (Lin et al., 2022; Wu et al., 2022; Fang et al., 2022) and used on end-to-end monomeric folding tasks, achieving approaching or equivalent accuracies to AlphaFold-based models. In this paper, we used the ESM-2 (Rives et al., 2021) model with 3 billion parameters as the base PLM.

### Further Pre-training

Based on existing explorations on PLMs, our work uses *further pre-training* to improve existing PLMs to capture inter-chain evolutionary patterns. Further pre-training is a widely used strategy in natural language processing for domain adaptation, where a pre-trained model is further trained to fit corpora in the target domain biased from the original one. Previous studies in Natural Language Processing (Lee et al., 2020; Beltagy et al., 2019; Sun et al., 2020) proved the effectiveness of further pre-training on domain transfer tasks, whereas the target domain generally has massive unlabeled data. In our case, notably, the known groups of interactive protein sequences are far sparser than monomeric ones.

## 3 Method

For an overall picture of our model, Figure 1 compares the workflows of protein complex prediction with i) MSA-based models such as AlphaFold-Multimer; ii) PLM-based (monomeric) models such as ESMFold and OmegaFold; and iii) our method Uni-Fold MuSSe+.

**Figure 1:**
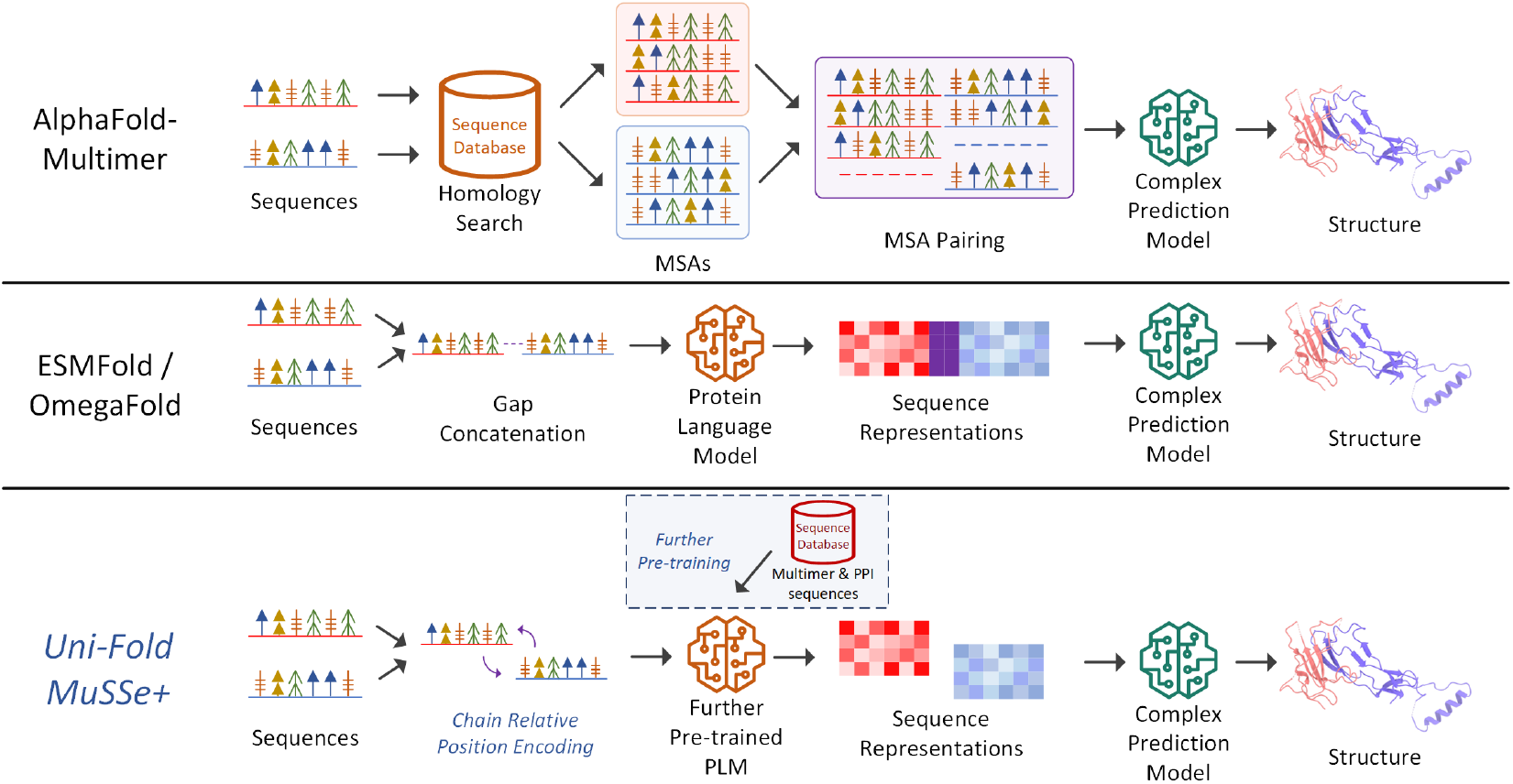
The workflows of MSA-based complex prediction methods, previous language modelbased methods such as ESMFold and OmegaFold and our proposed method Uni-Fold MuSSe+.

### 3.1 Further Pre-training of Protein Language Model

In order to further pre-train ESM-2, we carefully modified the architecture to adapt the model to inputs of both single sequence and multiple interactive sequences, without damaging the effects of the pre-trained parameters. To summarize, we introduce chain-level relative positional embedding to properly distinguish different chains, a cropping strategy for large protein complexes, and a data augmentation strategy to relieve overfitting.

#### Chain Relative Positional Encoding

Based on the Rotary Position Embedding (RoPE) (Su et al., 2021) used in ESM-2 (Lin et al., 2022), we propose two modifications to correctly introduce relative chain information. The first is to fix the incorrect RoPE among different chains. RoPE multiplies query and key embeddings in the attention module with sinusoidal embeddings so that the relative positional information can be captured by the Query-Key dot-product. However, although RoPE works well for single-sequence, it produces incorrect relative positional signals for residues from different chains. To address this issue, we proposed a fix for RoPE as follows: 1) First, a pair-wise matrix ***M*** is used to check if a residue pair is from the same chain (**M**_*ij*_ = 1) or not (**M**_*ij*_ = 0); 2) Besides the Query-Key dot-product with RoPE embedding, denoted as ***A***_RoPE_, we calculate an additional Query-Key dot-product without RoPE embedding, denoted as ***A***_raw_. 3) To remove the wrong signals of RoPE for different chains, we use the following update for the Query-Key dotproduct, ***A*** = ***A***_RoPE_ · ***M*** + (***A***_raw_ + Mean(***A***_RoPE_ — ***A***_raw_)) · (1 – ***M***), where Mean(·) is the mean pooling across the first two dimensions. This new update uses the RoPE-based attention for the residues on the same chain, and uses the non-RoPE attention, with the background deltas (average of differences), for the residues on the different chains.

##### Algorithm 1 Self-attention with chain relative position embedding

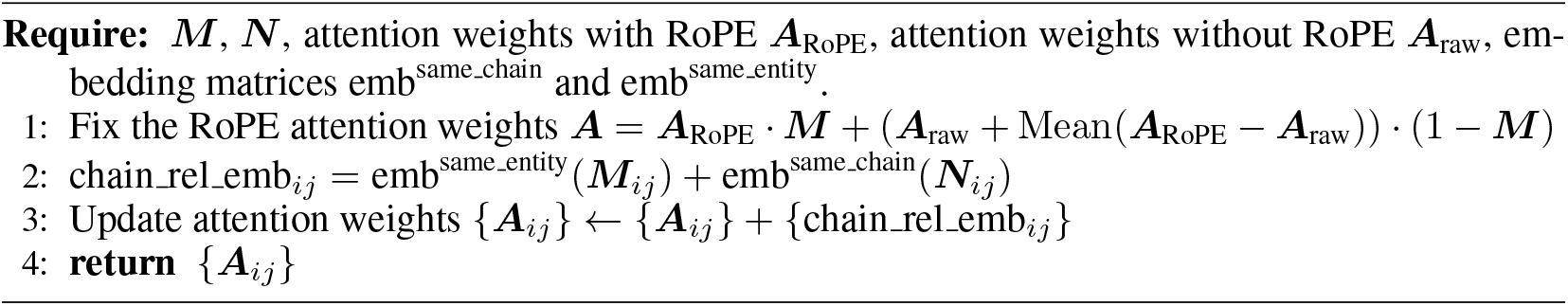

The second is to introduce the chain-level information. Besides corrected RoPE, we introduce two chain-level relative positional encodings. 1) ***M***, in which ***M***_*ij*_ encodes whether a residue pair is from the same chain or not. 2) ***N***, in which ***N***_*ij*_ encodes whether a residue pair is from chains of identical sequences or not, i.e., the possible different chains, but with the same sequence content. The two positional encodings are used as the index for embedding lookup tables and then are added to the attention weights before softmax in the attention module. Note that the parameters of embedding lookup tables are shared across layers. The details can be found in Algorithm 1.

#### Multi-chain Proportionally Contiguous Cropping

The total number of residues can be very large in multimeric sequences, which is infeasible to be fitted into GPU memory and inefficient in training. We use a cropping method similar to *ContiguousCropping* in AlphaFold-Multimer, with moderate changes, i.e., we crop chains proportionally according to their length. The details can be found in Algorithm 2.

##### Algorithm 2 Contiguous and Proportional Cropping

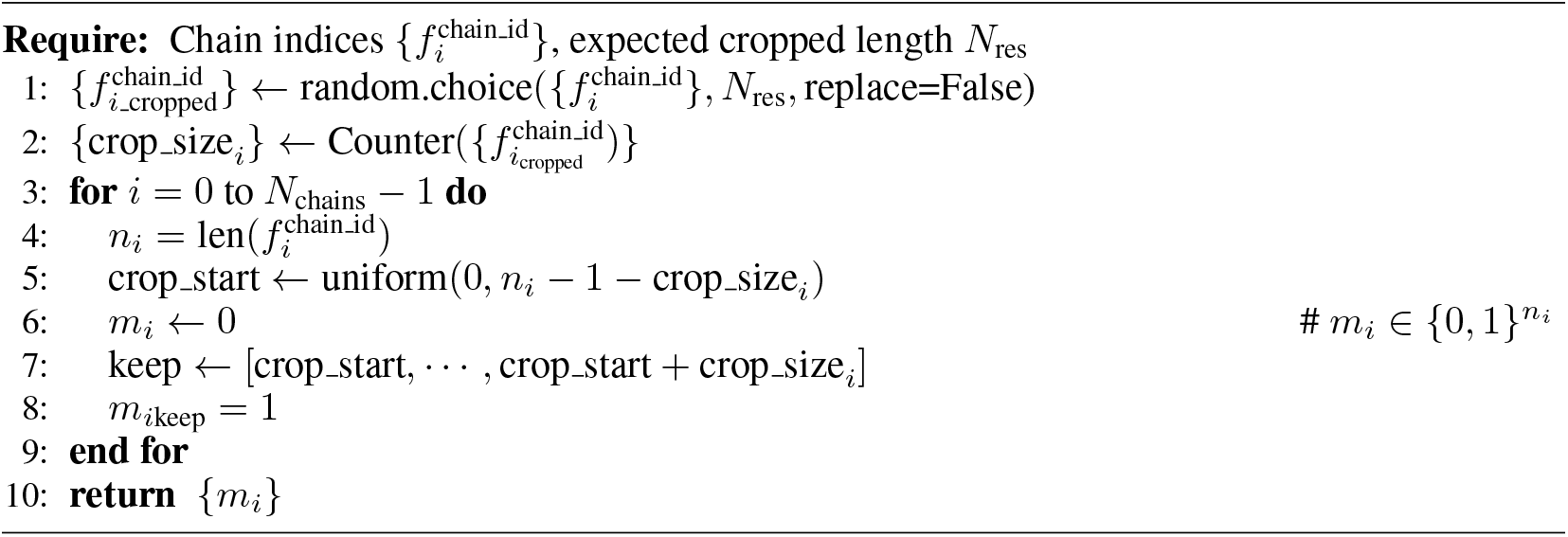

#### Data Augmentation

Directly using multimeric sequences of solved complex crystal structures for further pre-training PLMs can easily cause over-fitting, due to limited available data. Hence, we leverage more protein sequences from other sources, i.e., the sequence pair from protein-protein interaction data and the monomeric sequence. As the sizes of these datasets vary different, we assign different sample probabilities for them during training. The details of these datasets and the sampling probabilities for them are presented in Sec. 4.1.

### 3.2 Complex Prediction Model

We followed the major implementation of Uni-Fold Multimer (Li et al., 2022) to build the complex prediction model. The model consists of an encoder block which encodes the PLM representations, and a structure module which predicts the complex structure. The encoder block is adapted from the Evoformer (Jumper et al., 2021), simply by removing Extra-MSA blocks, template-related blocks, and the `MSAColumnAttention` block. The structure module is implemented as-is. We enlarged the embedding sizes of both the encoder and structure module to match the input dimensionality of ESM-2.

Similar to ESMFold (Lin et al., 2022), we use the token representations in the ESM-2 as the initial hidden states for structure prediction, with no gradient propagated back to the PLM. Notably, instead of using the linear combination of representations from all 48 layers, we simply combine those from the first and the last layers as inputs with learnable parameters *α, β*:

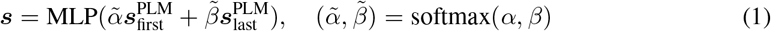

This is based on the observation that the first and the last layers contribute up to 97.3% to the folding model in the learned parameters of ESMFold (see the results in Figure 2). By doing so, the computational and storage costs are dramatically reduced. We used the relative chain and position encoding as the initial pairwise states.

**Figure 2:**
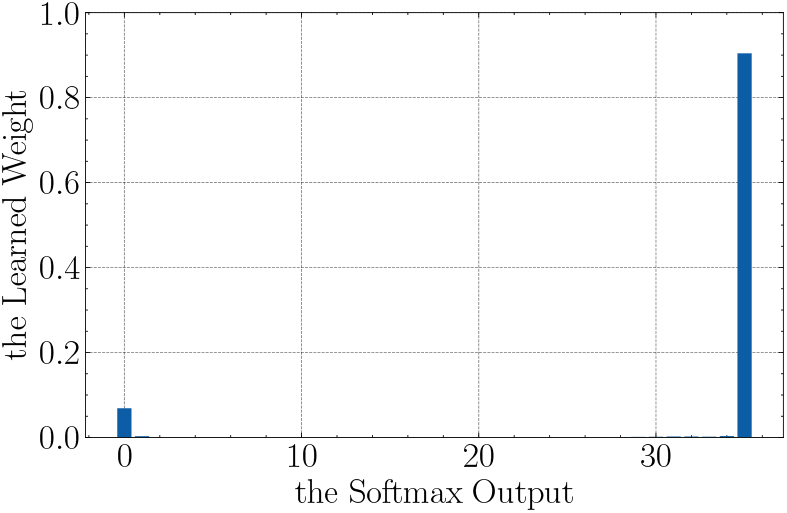
The probability distribution of learned weights for each layer output in ESMFold.

## 4 Experiments

### 4.1 Further pre-training of ESM-2

#### Model, Data and Training

We conducted further pre-training on ESM-2 (Lin et al., 2022), one of the largest pre-trained protein language model pre-trained on vast protein sequence data via masked language modeling. The model with 3 billion parameters is chosen as the base model.

A combination of three datasets is used in the further pre-training: i) `UniRef50`: approximately 50 million monomeric sequences in UniRef50 (Mirdita et al., 2017); ii) `PDB:` a total of 133 thousand groups of sequences of solved protein multimers in the Protein Data Bank; and iii) `PPI`: 2.6 million pairs of protein sequences known to have potential interactions from existing databases. Specifically for `PPI`, raw and partially overlapping interaction protein pairs were combined from seven sources: HINT (Das & Yu, 2012), intACT (Orchard et al., 2014), HIPPIE (Alanis-Lobato et al., 2016), prePPI (Zhang et al., 2012), BioGRID (Oughtred et al., 2019), comPPI (Veres et al., 2015) and huMAP (Drew et al., 2017). For higher data quality, we filtered out duplicated interactions, protein sequences outside of the Swiss-Prot database, and interactions with confidence scores lower than 0.8 in intACT, comPPI and huMAP. A total of 2,619,684 interaction pairs between 61,643 proteins were finally acquired.

The further pre-training included a total of 100,000 steps, each with a batch of 128 samples. Each sample contained one sequence in `UniRef50`, a group of multimeric sequences in `PDB`, or an interaction pair in `PPI`, with probability 2:4:4. We used AdamW (Loshchilov & Hutter, 2017) as the optimizer with *β*_1_. = 0.9, *β*_2_ = 0.98, and *L*_2_ weight decay of 0.01. The learning rate was linearly warmed up to 1*e*-5 in the first 2,000 steps, and linearly decayed to 5*e*-6 at the final step. This process was run on 32 NVIDIA A100 GPUs (80G) for approximately 30 hours.

#### Effectiveness of Further Pre-Training Strategies

We verified the effectiveness of the proposed method by the masked token prediction accuracy during further pre-training. In particular, we set up a baseline where the original ESM-2 model architecture was only with `PDB`. As is shown in Figure 3, the baseline was vulnerable to overfitting in further pre-training, whereas our method displayed better generalizabilty.

**Figure 3:**
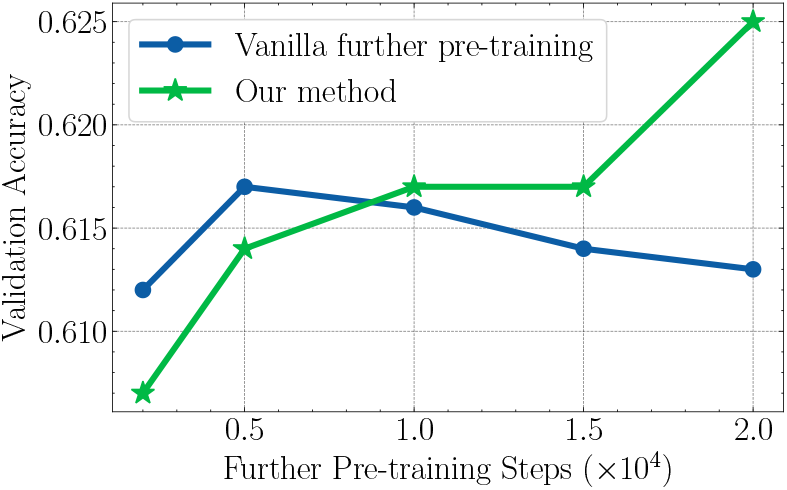
Mask token prediction accuracy on validation set during further pre-training.

### 4.2 Single Sequence Protein Complex Prediction

#### Dataset

For the training dataset, we follow the processing steps described in (Jumper et al., 2021) and (Evans et al., 2022). First, we use all the PDB complexes released before 2022-05-01, and then filter out all chains of which the resolution is greater than 9*Â* or there exists a residue whose occupancy frequency is greater than 0.8. For sequence clustering, we use MMseqs easy-cluster^1^ with default parameters at 40% sequence identity and result in 37812 sequence clusters, 133954 unique sequences, 645947 chains, and 179854 PDB assemblies. When training complex predictor, each PDB assembly *i* is sampled with unnormalized probability 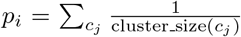, where *c_j_*
is the *j_th_* chain of assembly *i*, cluster_size(*c_j_*) is the size of the corresponding cluster of *c_j_*, and last the probability *p_i_* is normalized over all PDB assemblies. We also construct a distillation dataset with 914800 structure predicted by AlphaFold2, of which the mean pLDDT> 70. During training, we randomly sample distilled or PDB structures with probability 0.75 or 0.25, respectively. For inference data, we use all PDB complexes released after 2022-05-01, then filter out all multimeric data whose length-weighted sequence similarity is greater than 0.4 with any training sample, and randomly select one sample from each sequence cluster to constitute the last inference dataset.

#### Model and Training

We mainly followed the implementation of Uni-Fold Multimer (Li et al., 2022) to build the folding model. The hidden dimensions of Evoformer and the structure module were both extended to 1,024 with 32 attention heads. We used Adam (Kingma & Ba, 2014) *β*_1_ = 0.9, *β*_2_ = 0.999 as the optimizer. The learning rate was warmed up in the first 1,000 steps to a peak value of 0.001, and then decayed by 0.95 every 50,000 steps in stair-case style. The model was trained on 64 NVIDIA A100 GPUs (80G) with 150,000 steps and batch size 128 (by gradient accumulation) in about 15 days.

#### Baselines and Metric

We compared Uni-Fold MuSSe and MuSSe+ with i) AlphaFold-Multimer; ii) AlphaFold-Multimer with single sequence inputs, i.e. with MSAs and templates removed; iii) OmegaFold (Wu et al., 2022) and iv) ESMFold (Lin et al., 2022), in which we concatenated the sequences with 20-residue gaps embodied by the unknown symbol `X ([UNK])`. The only difference between MuSSe and MuSSe+ was that the latter used the further pre-trained PLM. Notably, both folding models in Uni-Fold MuSSe and MuSSe+ were trained from-scratch. Following Li et al. (2022), we used TM-score to evaluate the predicted complexes. Specifically, the optimal alignment between a prediction and the ground truth was calculated on the entire assembly structure. The scores were then calculated by averaging over all *C_α_* atoms in all chains. Best scores across all possible permutation alignments was reported.

### 4.3 Results

The results of the experiment are presented in Table 1. As can be seen, (1) AlphaFold-Multimer without multiple sequence alignment (MSA) lags behind OmegaFold, ESMFold, Uni-Fold MuSSe and Uni-Fold MuSSe+ by a considerable margin, highlighting the crucial role of language model representation in the absence of MSA information. (2) Uni-Fold MuSSe and Uni-Fold MuSSe+ outperform omegaFold and ESMFold, demonstrating that the incorporation of language model representations into the training process of protein complex prediction can further enhance performance. (3) Although Uni-Fold MuSSe and Uni-Fold MuSSe+ outperform OmegaFold and ESMFoLd, it still lags behind AlphaFold-Multimer with MSA input, indicating room for improvement in language model-based complex prediction methods. (4) Between the two variants of our method, although the average TM-Scores are similar, Uni-Fold MuSSe+ shows better performance in predicting high-quality complex structures, since it predicts more structures with TM-Score > 0.95 (and > 0.85). This is evident that Uni-Fold MuSSe+ is superior to the variant without further pre-training.

**Table 1:** Performance on multimeric protein structure prediction. The second and third columns are the mean and median TM-score among all test samples, and the right two columns are the percentage of predictions with a TM score above a predetermined threshold.

|  | TM-Score |  |  |  |
| --- | --- | --- | --- | --- |
|  | Mean | Median | % > 0.85 | % > 0.95 |
| AlphaFold-Multimer | **0.7106** | **0.9057** | **58.78** | **37.16** |
| AlphaFold-Multimer w/o MSA | 0.3012 | 0.2001 | 8.11 | 1.35 |
| OmegaFold (Wu et al., 2022) | 0.3986 | 0.2439 | 16.89 | 6.08 |
| ESMFold (Lin et al., 2022) | 0.4357 | 0.2715 | 22.97 | 7.43 |
| Uni-Fold MuSSe | **0.5871** | 0.6038 | 39.86 | 20.27 |
| Uni-Fold MuSSe+ | 0.5841 | **0.6675** | **41.22** | **28.38** |

We also assessed the performance of our method and baseline methods using the TM-Score and RMSD metrics at the sample level and present the results in Figure 4. In agreement with our previous conclusions, we also observed that Uni-Fold MuSSe+ produced a higher number of samples with TM scores close to 1.0 or 0.0, compared to Uni-Fold MuSSe without further pre-training. This observation offers valuable insights into how language models can be effectively utilized for protein structure prediction and serves as a basis for further improvement.

**Figure 4:**
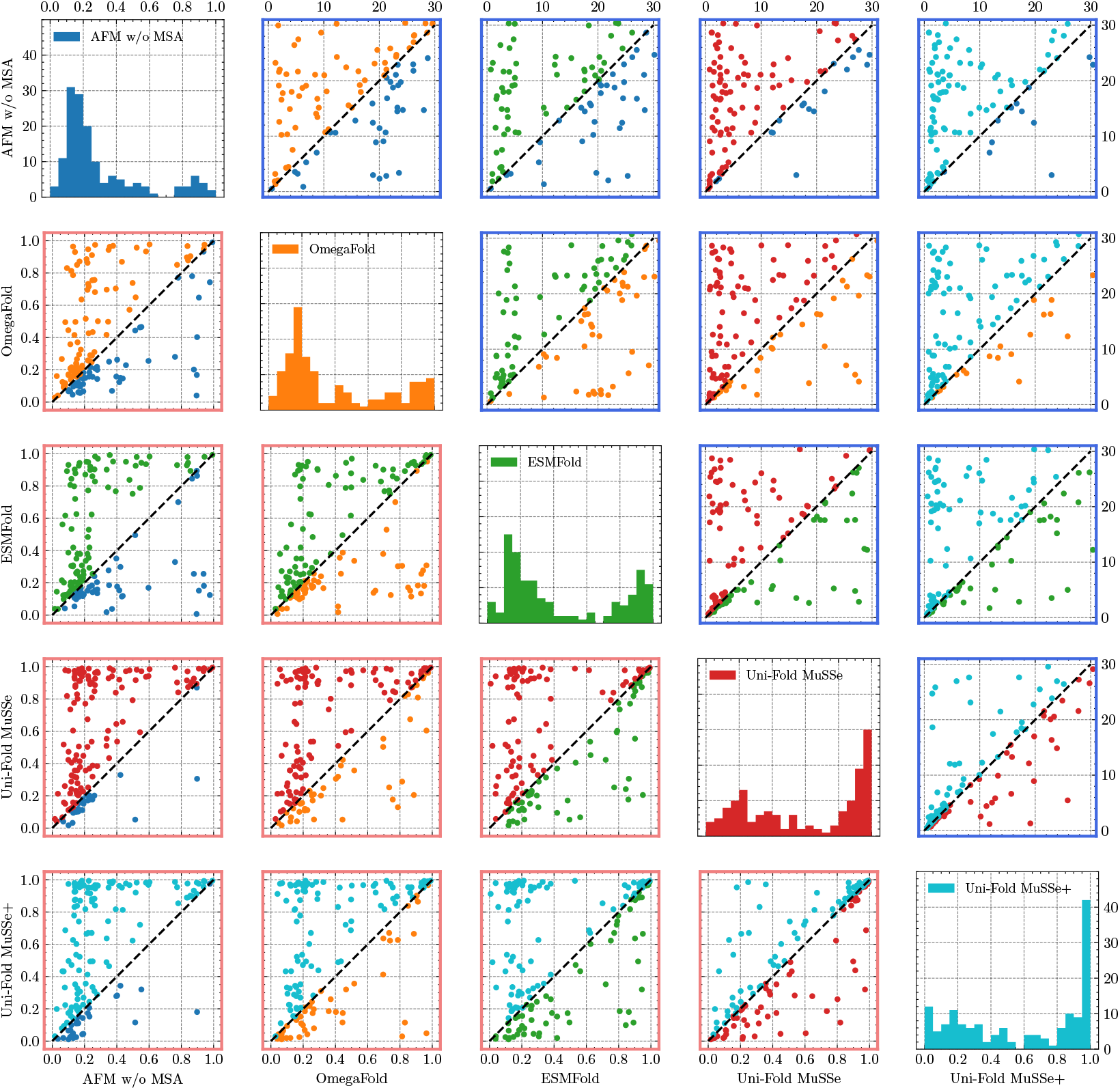
Scatter matrix of TM-score (bottom left, with red frames) and RMSD (top right, with blue frames) among our methods and baselines. Dots in different colors highlight that the corresponding method is better. The diagonal plots are histograms of TM-scores for each method.

Our analysis also discovered some instances on which Uni-Fold MuSSe+ outperformed AlphaFold-Multimer, which relies on homology search and pairing. Two cases can be seen in the comparisons presented in Figure 5, where the predictions of Uni-Fold MuSSe+ were found to be better to those of AlphaFold-Multimer. The limited number of cross-chain multiple sequence alignments (MSA) pairings for these complexes may explain this outcome, as seen in the cases of 7U5O with 348 pairings. These results highlight the potential of language-model based complex predictors in scenarios where the number of cross-chain MSA pairings is limited.

**Figure 5:**
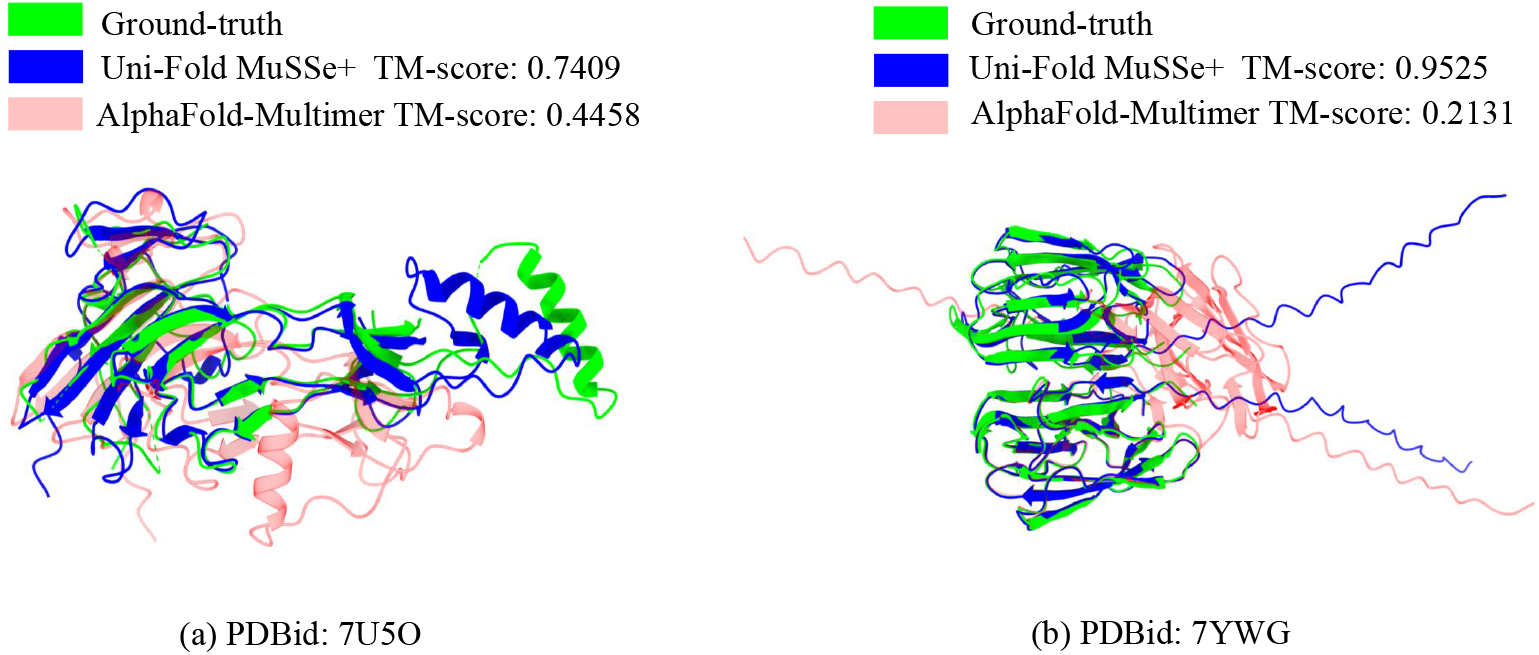
Predicted structures of two cases. Experimentally solved structures are colored in green, AlphaFold-Multimer predictions in red, and Uni-Fold MuSSe+ predictions in blue.

## 5 Conclusions

In this work, we proposed Uni-Fold MuSSe+, a complex structure predictor with no need for homology search and pairing. The core idea of Uni-Fold MuSSe+ is to resort to pretrained protein language model representation for evolutionary patterns rather than from multiple sequence alignment. We verify the effectiveness of our method on protein complex prediction, and can improve the baseline methods significantly. For future works, first, we will dig deeper into why the average performance of further pretrained language model is similar to the one without further pretraining. Second, further improving protein language model based structure predictor to the performance of MSA based works is a valuable direction. Finally, how to pre-train a multimeric language model efficiently and effectively is also an interesting topic.

## Acknowledgement

We would like to thank Xuyang Liu and Lüjun Guo for their contributions in preparing the UniRef50 and protein-protein interaction datasets for further pre-training.

## Footnotes

1 https://github.com/soedinglab/MMseqs2

